# B cells imprint adoptively transferred CD8^+^ T cells with enhanced tumor immunity

**DOI:** 10.1101/2021.03.16.435714

**Authors:** Aubrey S. Smith, Hannah M. Knochelmann, Megan M. Wyatt, Guillermo O. Rangel Rivera, Amalia M. Rivera-Reyes, Connor J. Dwyer, David M. Neskey, Mark P. Rubinstein, Bei Liu, Jessica E. Thaxton, Eric Bartee, Chrystal M. Paulos

## Abstract

Here we report a novel strategy to reverse the tolerant state of adoptively transferred CD8^+^ T cells against melanoma through *ex vivo* expansion with the TLR9 agonist CpG. T cells generated in the presence of CpG display potent anti-tumor efficacy without *in vivo* co-administration of high dose IL-2 or vaccination, which are classically required for effective treatment of solid tumors using adoptive cell therapies. CD8^+^ T cells adopt a unique proteomic signature and are characterized by an IL-2Rα^high^ICOS^high^CD39^low^ phenotype after CpG-mediated expansion. Surprisingly, we found that the presence of B cells, in the culture, was essential for imprinting CD8^+^ T cells with this phenotype and moreover purified B cells were sufficient to mediate the CpG-associated changes in T cells. These findings reveal a vital role for B cells in the generation of effective antitumor CD8^+^ T cells and have immediate implications for profoundly improving immunotherapy for patients.

**SUMMARY STATEMENT:** The TLR9 agonist CpG allows B cells to license adoptively transferred CD8+ T cells with potent tumor immunity. These licensed T cells have a unique proteomic signature, are marked by low CD39 and high ICOS and IL-2Rα expression, and engraft robustly *in vivo*.

## INTRODUCTION

Only a decade ago, the treatment options for patients diagnosed with late-stage solid malignancies were mostly ineffective and patient outcomes were bleak. With the advent of immunotherapies, including checkpoint blockade and adoptive cell transfer (ACT) therapy, a new era of cancer care is underway. Many patients with advanced cancer, like metastatic melanoma, experience objective responses, undergo long-term remissions and can sometimes be cured of their disease following delivery of T cell-based therapies^1^. However, as some patients receive ACT as a last resort, after progressing on multiple lines of therapy, and only 20% experience durable complete responses, it is paramount to find ways to improve cell therapy^1^. Critical to the success of ACT is host preconditioning with chemo- or radiotherapy. The preconditioning regimen promotes engraftment and antitumor activity of transferred T cells^2^. Beyond depletion of host immune cells that compete for homeostatic cytokines^3^ and/or suppress transferred cells (T regulatory cells (Tregs) or myeloid-derived suppressor cells (MDSCs))^4,5^, host preconditioning also alters gut microbiota homeostasis. The systemic release of gut microbes and microbial products, in turn, activates the innate immune system and augments the effectiveness of infused T cells via pathogen recognition receptors (PRRs)^6^.

Additional insights from the microbiome field have been used to improve immunotherapy, such as through the use of Toll-like receptor (TLR) ligands to induce immunity^7,8^. We and other groups have shown that TLR activation via synthetic microbial ligands improves ACT therapy when administered directly to mice^9,10^. While these findings are important, delivering TLR agonists alongside ACT therapy may induce toxic side-effects in patients and delivering TLR ligands *in vivo* may induce unwanted effects on other immune cells or even tumor cells^11^. Although TLR agonists have the potential to profoundly augment immunotherapy, the safest and most effective ways to use them for ACT remain unknown.

CpG DNA agonists of TLR9 have been widely used in both preclinical and clinical settings in combination with various therapies (e.g. vaccines, checkpoint blockade therapies, immune-agonistic antibodies, chemo/radiation) and as a monotherapy to induce antitumor responses (NCT03445533, NCT03618641, NCT03007732, NCT03831295, NCT03410901)^12–15,16–20^. Initially, several clinical trials reported activation of immune cells in patients with TLR9 agonist administration, but few individuals experienced a complete durable antitumor response^21^. Three recent trials, however, reported patient responses (PR and CR) to TLR9 agonists in combination with either low-dose local irradiation (IR) of a single tumor site, pembrolizumab, or ipilimumab^15,18,22^. Importantly, all three reports noted treatment-emergent adverse events (AEs) of grade 3 and/or 4 in a portion of the patients. In these and other reports, CpG is directly administered to the patient where the route and timing of treatment are likely to be important^23^. CpG is typically administered locally to tumor lesions, either subcutaneously or intratumorally, in clinical trials to promote targeted action to the tumor and avoid bioavailability issues that arise with systemic intravenous injection^23^. While local administration strategies such as these may be a valuable therapeutic option for some patients, they do somewhat limit the patient population to those with readily accessible tumor sites. Further, as severe AEs arise in many patients treated with combination therapies which include TLR9 agonist administration, determining the best way to exploit these agonists therapeutically while bypassing *in vivo* toxicity is paramount.

Herein, we developed a novel method in which CpG promotes efficacy of cell therapy without *in vivo* administration. We hypothesized that the efficacy of T cells could be improved by incorporating a TLR9 agonist into the *ex vivo* expansion protocol. When combined with cell therapy as part of the expansion process, CpG does not have any unwanted off-target effects *in vivo*. Moreover, this approach obviates the need to determine route and timing of CpG delivery. ACT with CD8^+^ T cells expanded with CpG was effective without *in vivo* IL-2 and vaccine adjuvants, which are typically necessary. To our surprise B cells played an essential role in imprinting CD8^+^ T cells with potent immunity. Overall, we describe how addition of CpG in culture propagates CD8^+^ T cells which display overt immunity against solid tumors *in vivo* and reveal mechanisms underlying their potency.

## RESULTS

### Potent antitumor T cells are generated with ex vivo CpG stimulation

We hypothesized that the TLR9 agonist, CpG, could be used *ex vivo* to augment T cell-based antitumor therapy. To test this idea, we employed the pmel-1 mouse model of ACT in which CD8^+^ T cells express a transgenic TCR that recognizes the gp100 epitope expressed by melanoma and healthy melanocytes^24^. On day 0, CpG or vehicle (endotoxin-free water) was added to whole pmel-1 splenocytes at the time of TCR stimulation with hgp100 peptide. The culture was expanded in the presence of IL-2 for one week to preferentially expand pmel-1 CD8^+^ T cells (henceforth referred to as pmel-1) to >95% purity. After one week of expansion, pmel-1 were infused into lymphodepleted mice bearing established B16F10 melanomas (**Fig. 1 A**). Pmel-1 expanded with CpG were remarkably therapeutic *in vivo*, as CpG-treated pmel-1 regressed melanoma in mice while Vehicle-treated pmel-1 treatment had little therapeutic impact (**Fig. 1 B**). Mice infused with CpG pmel-1 survived significantly longer than mice given vehicle pmel-1 or mice left untreated (**Fig. 1 C**). Moreover, pmel-1 engrafted and persisted at higher frequencies in the blood of mice if they were expanded in the presence of CpG *ex vivo* (**Fig. 1 D**). Of note, this robust response was achieved without co-administration of IL-2 or vaccine adjuvants *in vivo* — previously deemed necessary for durable immunity via pmel-1 ACT^24^. Thus, adoptively transferred CD8^+^ T cells can mediate remarkable responses to tumors and robustly persist *in vivo* when expanded *ex vivo* with CpG.

**Fig. 1.**
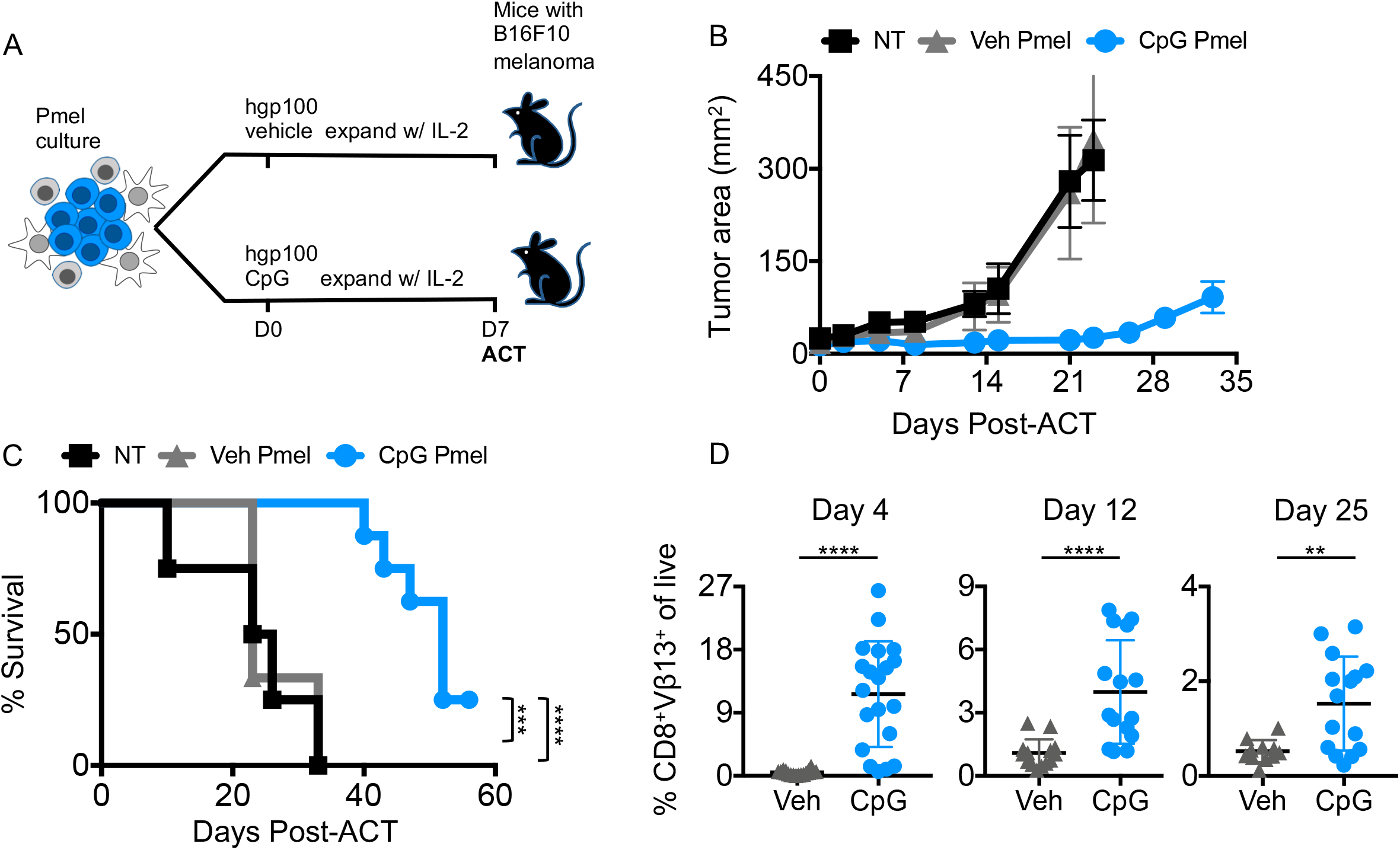
Potent anti-tumor T cells are generated with *ex vivo* CpG stimulation. A) Schema of experimental design. Pmel-1 splenocytes are activated with peptide +/− CpG on day 0 and expanded in IL-2 until D7 when they are assayed and infused into mice bearing 5-day established B16F10 melanoma. Mice were irradiated with 5Gy TBI 1 day prior to cell transfer. B) Tumor area over time of mice treated with vehicle pmel-1, CpG pmel-1 or NT. C) Survival of mice in B. D) Percentage of donor cells in peripheral blood of mice treated with vehicle pmel-1 or CpG pmel-1 on days 4, 12, and 25 post-ACT. Statistics: C) Log-rank test, D) Mann-Whitney test. **p< 0.01, ***p< 0.001, ****p< 0.0001.

### *In vitro* CpG stimulation generates T cells with a unique proteomic signature

As stark differences in antitumor ability were observed *in vivo,* we hypothesized that CpG treatment leads to the expansion of a CD8^+^ T cell product phenotypically distinct from traditionally expanded cells. To broadly assess how CD8^+^ T cells expanded from a CpG treated culture are different from traditionally expanded cells, we compared these T cell products using proteomic analysis. As portrayed in **Fig. 2 A**, proteins were extracted, purified and change in protein abundance from CpG treatment were determined using label-free LC-MS/MS-based proteomics of Day 7 CD8^+^ T cells. Not surprisingly, the proteomes of Vehicle and CpG-expanded CD8^+^ T cells were clearly divergent based on principal component analysis (PCA) (**Fig. 2 B**). Of over 2000 proteins identified between Vehicle and CpG-expanded T cells, there were 77 proteins differentially expressed between the two groups (Student t-Test with permutation-based FDR cutoff of 0.01 and S0 = 0.1) (**Fig. 2 C**). Several proteins involved in fatty acid metabolism (ACADLl, ACAD9, CLYBL) and GO Molecular Functions related to fatty acid metabolism were enriched in CpG-derived pmel-1 (**Fig. 2 C and D**). However, the most markedly increased protein in pmel-1 expanded with CpG was IL-2Rα (**Fig. 2C**). As IL-2Rα is required to form the high-affinity receptor for the T cell growth factor IL-2, this protein may be important for the antitumor function of CpG-generated T cells. Further, since IL-2Rα is expressed on the surface of T cells we posited that its expression level could be used to track the effects of CpG on expanded CD8^+^ T cells.

**Fig. 2.**
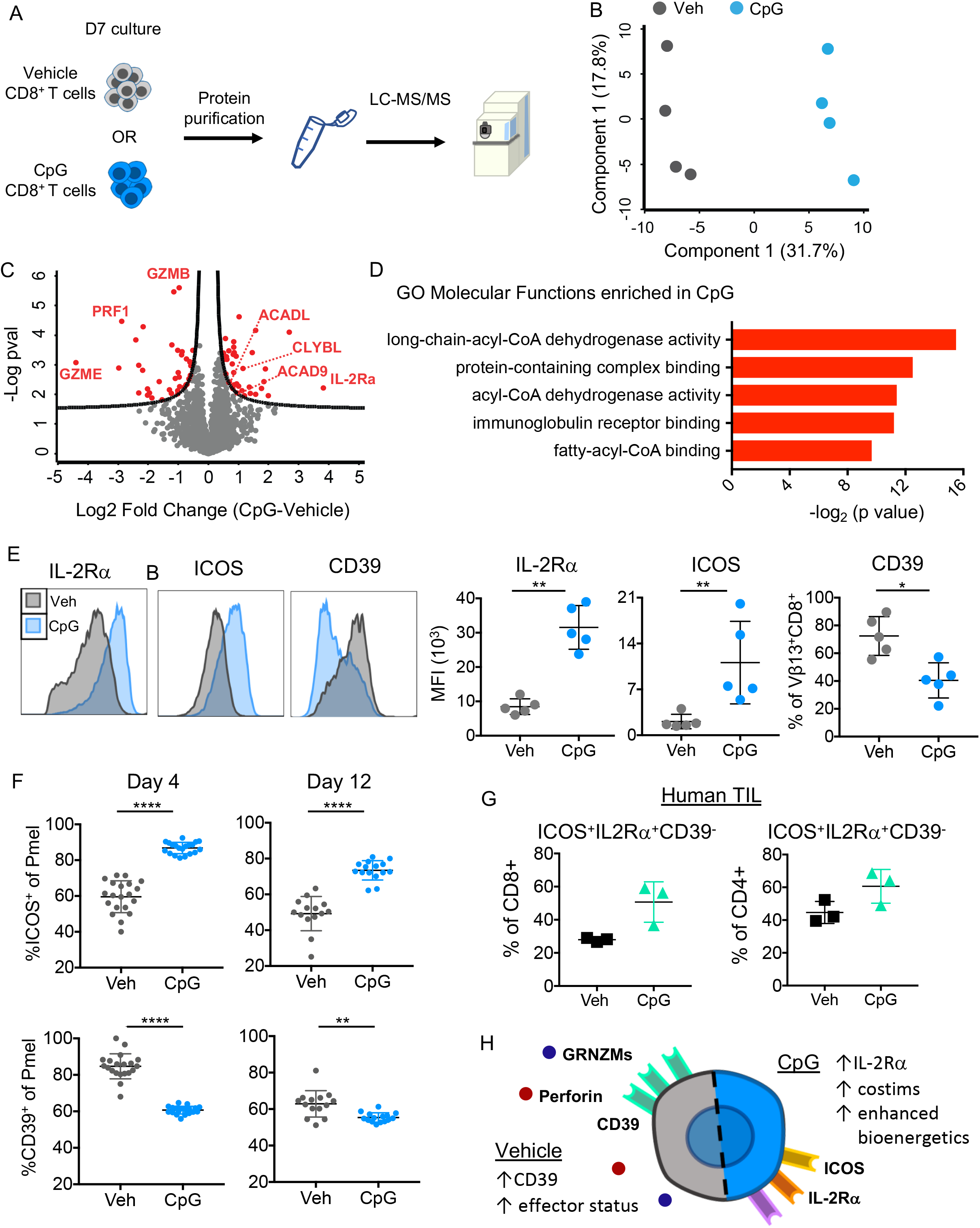
*In vitro* CpG stimulation generates T cells with a signature phenotype which is maintained post-ACT. A) Design of proteomic analysis of T cells; Vehicle or CpG-expanded T cells were collected on Day 7 of culture, subjected to protein extraction and purification, and then analyzed using LC-MS/MS. B) Principle component analysis of Vehicle or CpG-generated pmel-1 T cell proteomes. C) Volcano plot comparing protein expression between Vehicle and CpG-expanded T cells. The significance threshold was determined using a two-sided T test adjusted p value (FDR < 0.05) and an S0 parameter of 0.1. D) Top 5 GO: Molecular Functions enriched from proteins expressed more in CpG-generated pmel-1 T cells compared to Vehicle-generated cells (using https://toppgene.cchmc.org/enrichment.jsp). E) Representative flow cytometry histogram (left) and replicates (right) of extracellular expression markers from D7 of cell culture of vehicle or CpG treated pmel-1. F) Percentage of donor pmel-1 cells in the blood expressing ICOS (top) or CD39 (bottom) on D4 or D12 post ACT. G) Percentage of human HNSCC TIL expressing IL-2Ra and ICOS, but not CD39 after expansion with CpG. H) Diagram of CpG-generated T cell characteristics. Statistics: C) Significance threshold was set using two sided t test adjusted p-value (<0.05 FDR) and a S0 parameter of 0.1, E-F) Mann-Whitney U test. ns, not significant, *p< 0.05, **p< 0.01, ***p< 0.001, ****p< 0.0001.

We used flow cytometry to confirm the heightened expression of IL-2Rα and further investigate the T cell phenotype of CpG pmel-1. In agreement with our proteomics analysis, CpG-expanded pmel-1 expressed heightened IL-2Rα (**Fig. 2 E**). In addition, we found the costimulatory molecule ICOS was expressed to a higher degree on the surface of CpG-generated CD8+ T cells. IL-2Rα and ICOS have been reported to augment T cell engraftment and function^25–27^. CpG pmel-1 also expressed less CD39, an immunosuppressive ectoenzyme expressed on exhausted CD8^+^ T cells^28-29^, than vehicle pmel-1. Thus, when expanded in the presence of CpG, pmel-1 cells display a signature phenotype (IL-2Rα^high^ICOS^high^CD39^low^) (**Fig. 2 E**). Note, in experiments henceforth, we tracked and used this signature profile as a metric to predict the effect of CpG on pmel-1 CD8^+^ T cells. Importantly, expression of the signature markers is maintained post ACT as heightened ICOS and diminished CD39 remained characteristic of the CpG-treated donor cells persisting in the peripheral blood (**Fig. 2 F**).

To determine if this phenotype would be recapitulated in human cells we cultured TIL from a patient with oral cavity squamous cell carcinoma with human CpG or vehicle at the start of TIL seeding in culture. Like their mouse counterparts, human T cells, both CD4s and CD8s, had larger proportions of ICOS^+^IL2Rα^+^CD39^-^ T cells post *ex vivo* culture with CpG (**Fig. 2 G**). Thus, both mouse and human T cells gain these desired surface markers after CpG treatment *ex vivo.* While the functional significance of this signature phenotype on CpG-generated T cell products remained incompletely elucidated, individually each marker has been associated with more or less fit T cells. Thus, we turned our attention to uncovering how merely adding CpG to cell cultures could bolster T cell potency.

### CpG treatment indirectly alters the CD8^+^ T cell phenotype and improves tumor immunity

We first questioned whether CpG was directly acting on pmel-1 T cells or if CpG activates other immune cells present at the onset of the culture, imprinting pmel-1 T cells with the signature phenotype and antitumor ability. Using the online database, ImmGen, we queried *TLR9* expression via available RNAseq datasets and found that many B cell, DC, and myeloid subsets express *TLR9* transcripts. Conversely, T cell subsets, from both healthy mice and those recently exposed to virus (7 days post-LCMV), express trace or undetectable *TLR9* transcripts in the mouse (**Fig. S1 A)**. In our mouse model, we corroborated these findings; TLR9 is not expressed in pmel-1 CD8^+^ T cells but is expressed by professional antigen-presenting cells (APCs), which are present only at the start of the TCR-stimulated culture (**Fig. S1 B**). Thus, pmel-1 likely acquire their unique phenotype (IL-2Rα^high^ICOS^high^CD39^low^) after CpG conditioning via TLR9-mediated activation of APCs. The TLR9-expressing APCs in the spleen include B cells, macrophages, and dendritic cells, and are present only at the start of culture. These APCs rapidly diminish while CD8^+^ T cells preferentially expand due to peptide-mediated triggering of the pmel-1 TCR. Specifically, few APCs are detected three days after adding peptide to the pmel-1 splenocyte culture (not shown).

Using this kinetic information, we designed an experiment to determine whether CpG can act directly on the T cells to confer the signature phenotype and improved tumor immunity. CpG was added at the start of culture on day 0 (Early CpG) when APCs and T cells are present, or CpG was added on Day 3 (Late CpG) when pmel-1 CD8^+^ T cells dominated the majority of the culture (**Fig. 3A**). Pmel-1 CD8^+^ T cells were assayed for surface IL-2Rα, ICOS, and CD39 one week after expansion. As anticipated, pmel-1 treated with CpG early expressed high IL-2Rα and ICOS and low CD39 one week after expansion (**Fig. 3 B**). Conversely, when CpG was added on day 3 of culture when nominal APCs remained, Late CpG pmel-1 expressed these surface markers to the same extent as vehicle-treated pmel-1 (**Fig. 3 B**). Early CpG pmel-1 persisted *in vivo* and improved therapeutic outcomes and survival (**Fig. 3, C-F**). However, Late CpG pmel-1 failed to persist in the blood or tumor of mice and did not improve survival (**Figure 3, D-F**).

**Fig 3.**
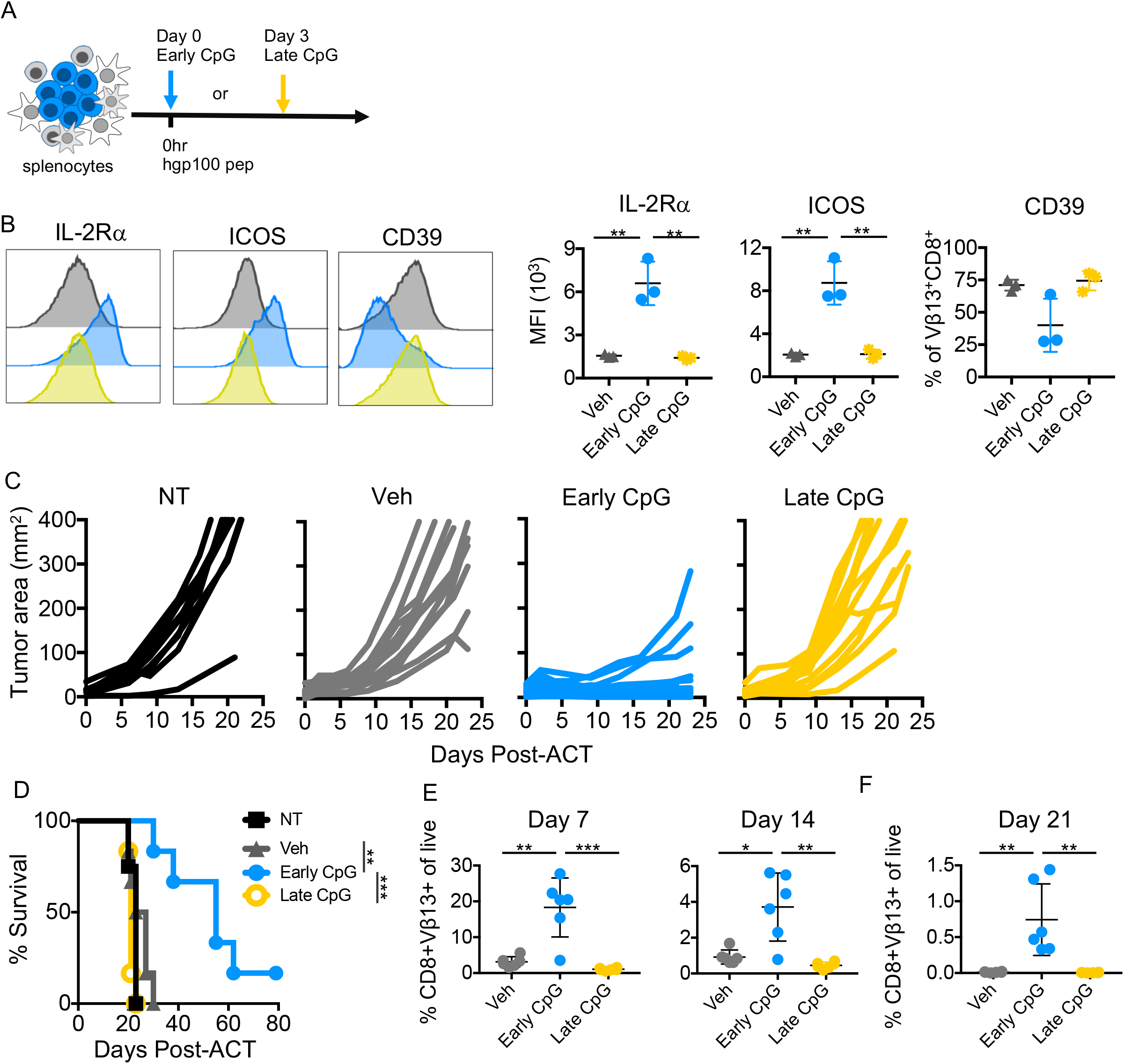
CpG treatment indirectly alters the CD8^+^ T cell phenotype and improves tumor immunity. A) Schema of experimental design; pmel-1 splenocytes were treated with CpG at the time of activation (Early) or 3 days post activation (Late) with peptide or treated with vehicle at the time of activation. B) Phenotypic marker expression on D7 of culture. C) Tumor area over time (single curves represent 1 mouse) treated with NT, vehicle pmel-1, Early CpG pmel-1, or Late CpG pmel-1. D) Survival of mice in C. E) Percentage of donor cells in the blood of mice from C on D7 and D14 post ACT. F) Percentage of donor cells in the tumor of mice treated in C on D21 post ACT. Statistics: B) Two-sample T-test, D) Log-rank test, E,F) Mann-Whitney U test *p< 0.05, **p< 0.01, ***p< 0.001, ****p< 0.0001.

Our data suggested that either the APCs present at the start of culture or the presence of CpG at priming were critical to the effects it imparted to antitumor CD8^+^ T cells. Therefore, we performed another experiment to determine if CpG could act directly on pmel-1 in the absence of APCs. Purified pmel-1 CD8^+^ T cells (negative isolation, >90%) were activated and expanded in the absence or presence of CpG; cells expanded logarithmically as expected **(Fig. S2 A)**. As anticipated, purified pmel-1 CD8^+^ T cells did not gain the signature phenotype or enhanced antitumor efficacy when expanded in the presence of CpG (**Fig. S2, B and C**). Moreover, the engraftment and persistence of enriched pmel-1 T cells treated with CpG were poor (**Fig. S2 D**). Taken together, these findings indicate that CpG does not modulate T cell phenotype directly. Instead, we reasoned that the CpG-mediated effects occur via other immune cells that express TLR9.

### The phenotype of CpG-generated CD8+ T cells is largely dependent on B cells

While many different immune cells are present in the early stages of CD8^+^ TIL cultures, intracellular TLR9 is expression is limited to B cells, DCs, and macrophages in the mouse (**Fig. S1 B**). Thus, we next sought to identify which APC was important for generating pmel-1 with the signature phenotype associated with CpG-promoted tumor immunity. We hypothesized that dendritic cells (DCs) were critical, as they are potent TLR9^+^ APCs that secrete cytokines and express heightened MHC I and II molecules when activated with CpG^30–33^. To address this question B cells, DCs, or macrophages were depleted from the starting cell culture prior to expansion for 7 days and phenotypic analysis. As additional controls, we also removed CD4^+^ T cells or NK cells before the start of culture as these populations are also present in pmel-1 splenocytes. In contrast to our hypothesis, depleting DCs from the CpG treated cell culture did not hinder the outgrowth of pmel-1 with the signature phenotype. Instead, the only depletion that prevented expansion of IL2Rα^high^ICOS^high^CD39^low^ pmel-1 was depletion of B cells (**Fig. 4 A**). Indeed, whether macrophages, CD4^+^ T cells, or NK cells were depleted, CpG treatment could still propagate IL2Rα^high^ICOS^high^CD39^low^ pmel-1 cells (**Fig. 4 A and Fig. S3 B**).

**Fig. 4.**
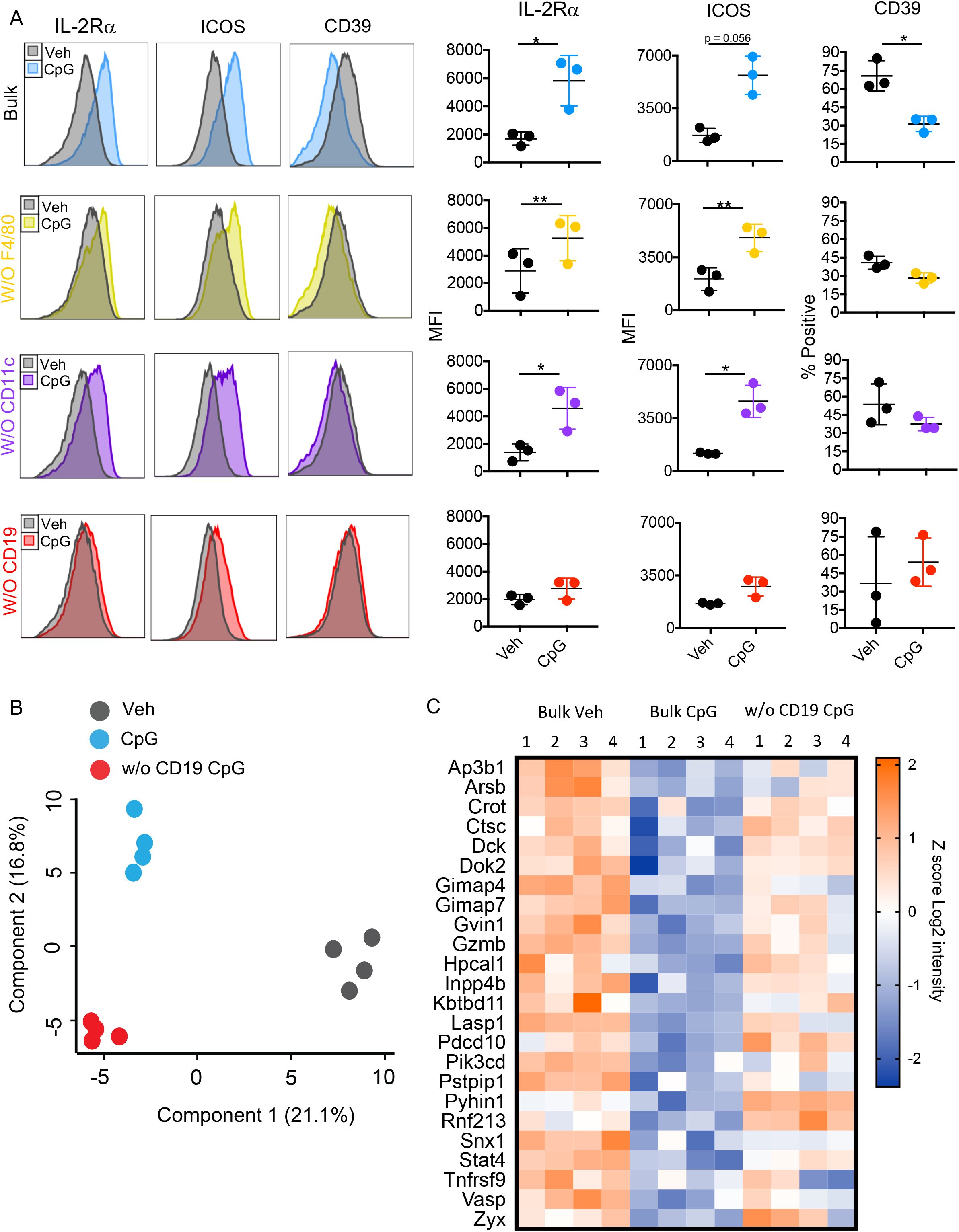
The phenotype of CpG-generated CD8+ T cells is largely dependent on B cells. A) Histograms (left) and biological replicates (right) of the expression of signature phenotype markers on the cell surface on Day 7. T cells were expanded from Bulk, F4/80-depleted, CD11c-depleted, or CD19-depleted cell cultures treated with Vehicle control or CpG. B) Principle component analysis of proteomic signatures of Day 7 expanded T cells derived from a Bulk Vehicle or CpG-treated culture, or a CD19-depleted CpG-treated culture. C) Heatmap comparing Z scores (of the Log2 protein intensity value) from the T cells expanded from the same groups as in B. Displayed are proteins that were expressed significantly higher (determined using two-sided T test adjusted p value (FDR < 0.05) and an S0 parameter of 0.1) in Bulk Vehicle generated T cells compared to Bulk CpG generated T cells. Statistics: A) Unpaired T tests. ns, not significant, *p< 0.05, **p< 0.01. Abbreviations: SCR = scramble ODN

Though the signature phenotype of CpG-expanded pmel-1 was preserved in all but the B cell depleted cultures, this analysis represents more of a snapshot of the likely myriad changes that occur with CpG expansion and which may be lost with the removal of B cells. Thus, we used proteomics to determine on a larger scale the characteristics of T cells expanded with CpG when B cells were present or absent. We included traditionally-expanded T cells (Bulk Vehicle) as a control to see whether removing B cells essentially reverts the phenotypic effects of CpG to that of a Vehicle-derived population. By PCA none of the ACT products (Vehicle, CpG-expanded, or CpG-expanded CD19 depletion) overlapped indicating that each of these groups was different from one another at the proteomic level (**Fig. 4 B**). We next tested if depleting B cells results in the expansion of a cell product that more closely resembles the ineffective Vehicle-derived CD8^+^ T cells. We found that a majority of the proteins which were significantly enriched (Student t-Test with permutation-based FDR cutoff of 0.01 and S0 = 0.1) in Vehicle-expanded pmel-1 were also more abundant in CD19-depleted CpG treated cells compared to T cells expanded from a Bulk culture treated with CpG (**Fig. 4 C**). Under certain circumstances, several of these proteins that were abundant in both the Bulk Vehicle and CD19-depleted CpG expanded pmel-1 have been associated with highly activated or terminally exhausted T cells including: GZMB, TNFRSF9, and PIK3CD^34,35^. However, these same proteins can also promote activation, costimulation, and cytotoxic function of T cells. These results indicate that CD19^+^ B cells, in the context of CpG, drive broad phenotypic alterations in T cell products, but do not reveal whether B cells are critical for the potent tumor immunity seen after expansion with CpG.

### Removal of B cells from the culture ablates the improved antitumor efficacy with CpG

Removing B cells from the start of culture was the only APC depletion that hindered the outgrowth of pmel-1 with the signature phenotype (IL2Rα^high^ICOS^high^CD39^low^), as shown in Figure 4. Moreover, if CpG was added to cell culture but B cells were depleted, the resultant cell product was phenotypically similar to the ineffective Vehicle-expanded T cells. In light of these results, we hypothesized that B cells would be similarly critical in conferring the antitumor ability of CpG-expanded pmel-1. To test this idea, we depleted B cells (CD19^+^ cells) from the culture as above prior to expansion and ACT. We also performed additional cell depletions from the starting culture to test whether other APCs (CD11c^+^ DCs or F480^+^ macrophages) or cell populations (CD4^+^ T cells or NK1.1^+^ NK cells) were necessary for the potent cell therapy seen with CpG. These cultures were then peptide activated and treated with or without CpG. In agreement with our hypothesis, we found that when B cells were removed from the culture, CpG pmel-1 cells were as ineffective as vehicle pmel-1 cells at regressing tumors *in vivo* (**Fig. 5 A**). Conversely, when DCs, macrophages, CD4^+^ T cells, or NK cells were removed from the CpG treated culture, the resultant expanded pmel-1 remained highly effective against melanoma when infused into mice and extended their survival (**Fig. 5 A and Fig. S3 C**). No survival advantage was achieved in mice treated with CpG pmel-1 that were expanded in the absence of B cells *ex vivo* (**Fig. 5 B**). Engraftment and persistence of pmel-1 in the blood were improved if cultures were treated with CpG, even in the absence of DCs, macrophages, NK or CD4^+^ T cells, at the start of culture (**Fig. 5 C and Fig. S3 E**). In contrast, the engraftment and persistence of CpG-treated pmel-1 cultures depleted of B cells were poor (**Fig. 5 C**). These findings uncover a key role for TLR9-activated B cells in generating CD8^+^ T cells with potent antitumor immunity.

**Fig. 5.**
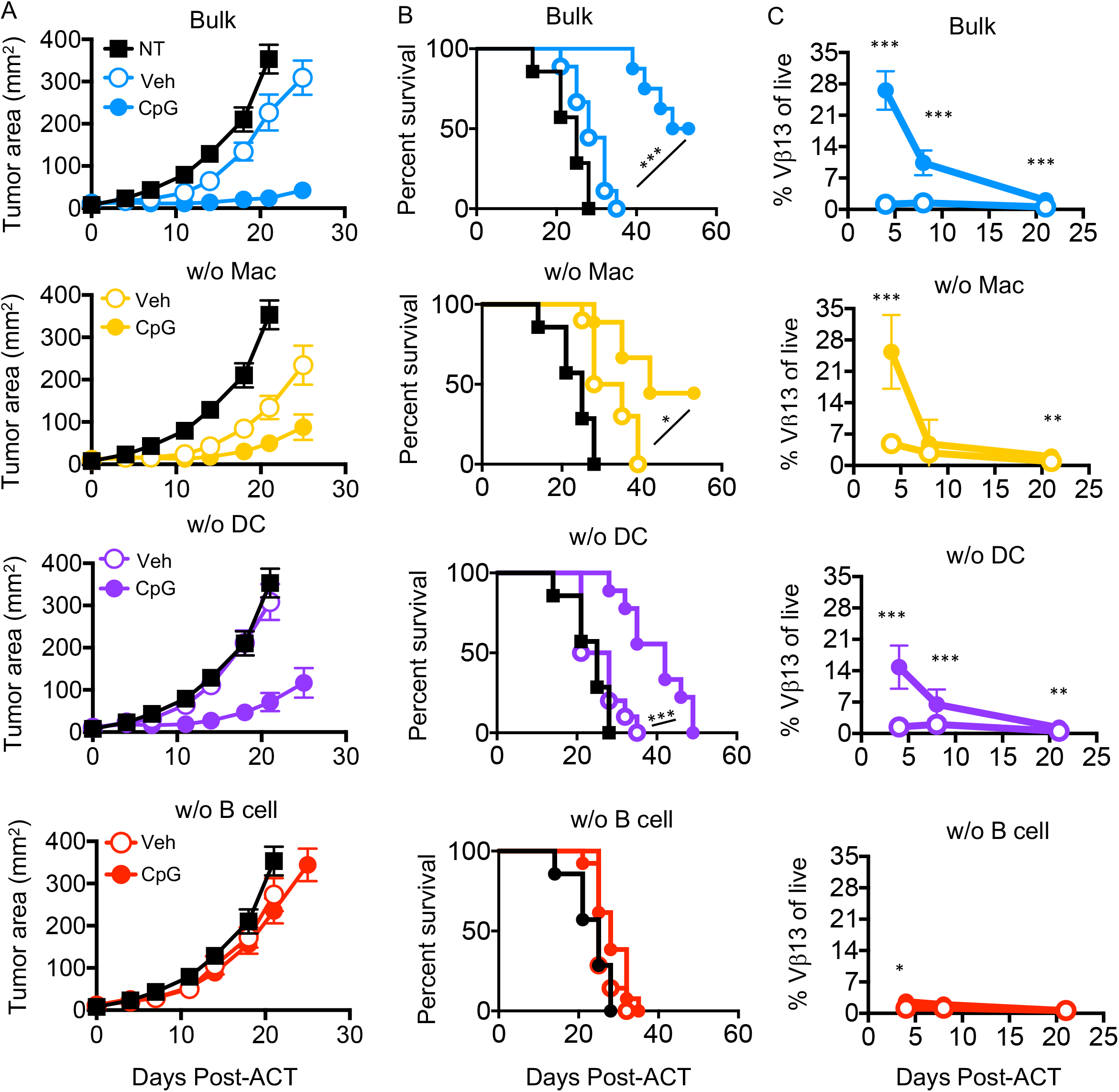
Removal of B cells from the culture ablates improved antitumor efficacy with CpG. A) Tumor area over time of mice treated with NT, or Veh or CpG-generated T cell products from Bulk, macrophage, DC, or B cell-depleted cultures. B) Survival of mice in A. C) Percentage of donor cells in the blood of mice treated in A on D4, D8, and D21 post-ACT. Statistics: B) Log-rank test. C) Mann-Whitney U at each time point. ns, not significant, *p< 0.05, **p< 0.01, ***p< 0.001, ****p< 0.0001.

### B cells alone are sufficient to improve antitumor T cells after activation via TLR9

Based on our previous data, we assumed purified B cells could enhance pmel-1 CD8^+^ T cells for ACT via TLR9 activation via CpG, but it remained unknown if the same extent of tumor immunity would be mediated if B cells were the sole APC. Thus, we examined the signature phenotype and antitumor properties of CpG-generated cells in bulk and purified B cell/T cell co-cultures. As seen in bulk cultures treated with CpG, when CD8^+^ T cells were expanded from a co-culture of CD8^+^ T cells and B cells treated with CpG, T cells also expressed more IL2Rα and ICOS. CD39 expression was similar between both CpG and vehicle treated groups in the purified B plus T cell culture, but the overall expression of both groups was lower than that of vehicle T cells generated from a bulk culture (**Fig. 6 A and Fig. S4 A).** Mice given either of the CpG-derived pmel-1 products arrested tumor outgrowth and enhanced survival time compared to mice treated with vehicle cells or those that were not treated (**Fig. 6 B-D)**. Survival did not differ between the two CpG treatment groups, suggesting that B cells induce the potency of pmel-1 T cells via CpG. Of note, pmel-1 generated without CpG, but from a B and T cell co-culture, were more effective antitumor T cells than their bulk vehicle counterparts, as demonstrated by both the slower outgrowth of tumors and longer survival (**Fig. 6 B-D**). Also, mice treated with either CpG-generated cell product had more donor pmel-1 cells in their blood one week after infusion, indicating better engraftment of CpG-generated cells. Though donor cells were more abundant in mice given the bulk CpG-treated product over those given the pure B/T cell CpG-treated product (**Fig. 6 E**), this trend was not observed for survival or tumor growth (**Fig. 6 B-D**).

**Fig. 6.**
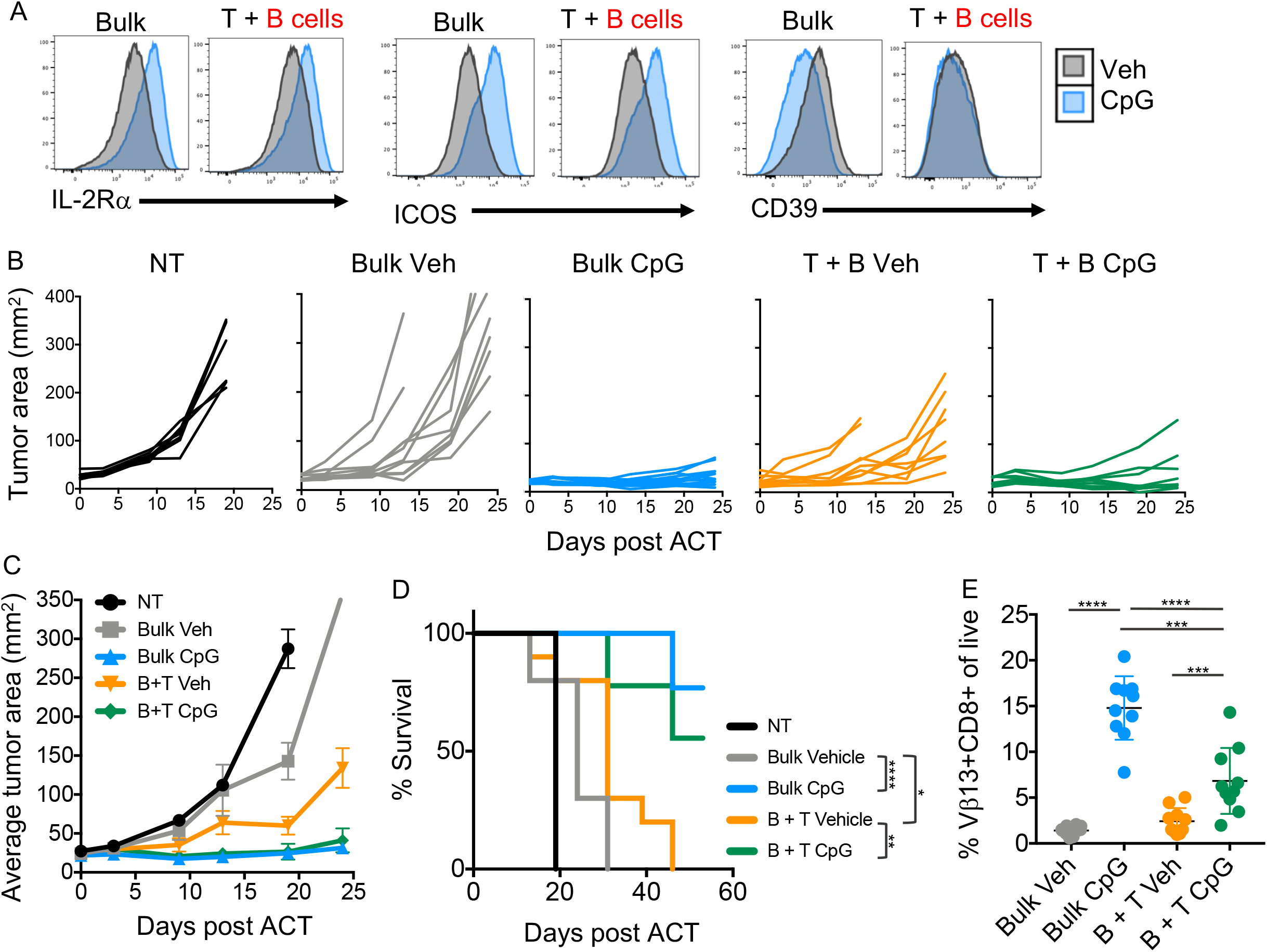
B cells alone are sufficient to improve antitumor T cells after activation via TLR9. A) Phenotype of day 7 T cells. B) Tumor area over time (single curves represent 1 mouse) of mice treated with NT, Bulk vehicle pmel-1, Bulk CpG pmel-1, B +T vehicle pmel-1, or B + T CpG pmel-1 (bulk indicates the T cell product was grown out of a whole splenocyte mixture seeded on day 0 while B + T indicates the T cell infusate was derived from a culture of purified B cells co-cultured with purified T cells seeded on day 0). C) Average tumor area over time of mice treated in B. D) Survival of mice in B. E) Percentage of donor cells (Vb13+CD8+) in peripheral blood of mice treated with each group listed in B on day 7 post-ACT. Statistics: D) Log-rank test, E) Mann-Whitney U test, *p< 0.05, **p< 0.01, ***p< 0.001, ****p< 0.0001.

Collectively, we found that a highly effective CD8^+^ T cell product can be achieved by simply adding the TLR9 agonist, CpG, to cell expansion protocols. Moreover, the mechanism of action of CpG is mediated by B cells, and B cells alone are sufficient to promote this powerful antitumor cell product via CpG, as depicted in summary figure (**Fig. 7**).

**Fig. 7.**
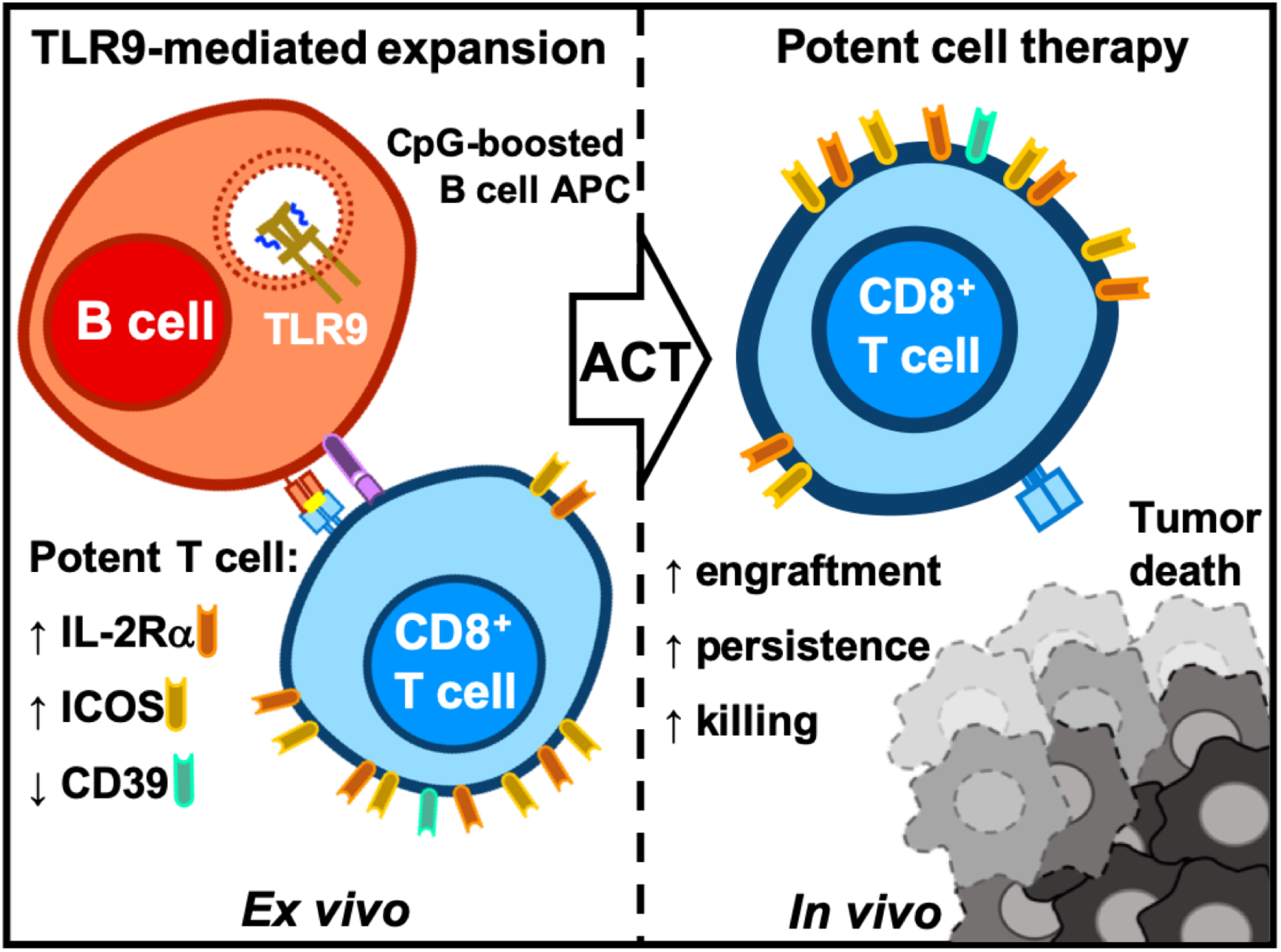
CD8+ T cells break tolerance to tumors in a B cell-dependent manner via TLR9 signaling. During the expansion of tumor-reactive T cells (left) CpG can be added to cell culture to improve the potency of cell therapy *in vivo* (right). *Ex vivo* B cells are responsible for improving the T cell product via CpG and T cells gain a unique phenotype: IL-2Rα^high^ICOS^high^CD39^low^. Once transferred CpG-generated T cells display enhanced engraftment, persistence, and tumor killing compared to a traditionally-expanded cell product.

## DISCUSSION

Herein, we describe a novel use for a long-studied microbial ligand, the synthetic Toll-like receptor 9 agonist CpG DNA. We reveal CpG can be added to the *ex vivo* cell expansion process to reverse the tolerant state of CD8^+^ T cells to tumors *in vivo* and augment cancer treatment. Perhaps most surprising, B cells were essential to the manifestation of this potent T cell product as observed through our cell depletion and purified co-culture studies.

The properties which drive CpG-generated T cells to robust antitumor responses *in vivo* are still unknown. However, our proteomic screen of T cell products points to increased fatty acid oxidation (FAO) in CpG-expanded cultures compared to traditionally-expanded T cells. Several reports demonstrate FAO is important for the development of memory T cells and their enhanced survival^36–38^. As CpG-derived T cells survive markedly longer *in vivo* compared to Vehicle-derived T cells, this advantage may in part be due to FAO-based energetics, however, further investigation is required to determine if FAO plays a role in the effectiveness of this T cell product. While some proteins of this metabolic pathway were enriched, IL-2Rα stood out as the most dramatically enriched protein in CpG-generated T cells. We confirmed this finding with flow cytometry and were also able to identify an expression pattern of two other markers of CpG-generated T cells – increased ICOS and decreased CD39.

Given our finding that CpG-induced T cells increase IL-2Rα and ICOS expression but express less CD39, a natural question follows: Do these molecules influence the potency of anti-tumor T cells? Our team previously reported that IL-2Rα on activated T cells allows them to respond in an improved and temporal fashion to IL-2, *in vivo*, and mediate anti-tumor immunity^*25*^. Further overexpression of IL-2Rα improves T cell tumor control^39^. ICOS, which is also highly elevated on CpG-treated pmel-1 T cells, supports the function and antitumor activity of various types of cell products (TILs, CARs) ^26,27^. ICOS signaling on T cells is also crucial for an anti-tumor response post-CTLA4 blockade^40^. Finally, CpG-mediated downregulation of the ectoenzyme CD39 may improve T cell tumor immunity. As CD39 is the rate limiting enzyme in the metabolism of pro-inflammatory ATP to immunosuppressive adenosine, a diminishment of CD39 may lead to less inhibitory adenosine within the tumor^41^. Moreover, CD39 marks effector T cells that have an exhausted phenotype in patients with infectious disease and cancer and have reduced survival after activation in older individuals^28,29 42^. In patients with metastatic melanoma, Krishna et al. demonstrated that CD39 and CD69 expression can be used to bifurcate patient response to cell therapy and that some CD39^-^ TIL from these patients are neoantigen recognizing stem-like T cells, which are known to potentiate powerful antitumor responses^43^. Thus, it is likely that the reduced CD39 and overt expression of IL-2Rα and ICOS on T cells post CpG treatment potentiate T cell function, metabolism, and persistence *in vivo*. The contribution of these surface molecules and others on the efficacy of this therapy will be explored in future studies.

Using two experimental approaches – one in which CpG was added to purified T cells and one in which CpG was added when few APCs remained in culture - we discovered that CpG does not directly alter CD8^+^ T cell phenotype or antitumor ability. This finding was not surprising as CD8^+^ T cells express nominal TLR9. Instead, our data showed that B cells in the CpG-treated culture were vital for enhancing T cells for ACT. CpG-generated cells remained effective against cancer if DCs, macrophages, CD4^+^ T cells, or NK cells were removed from the culture; however, removing B cells completely negated the CpG-improved cell therapy. Interestingly, we noted that not all of the cell products from these depletion strategies were equivalent in tumor efficacy compared to their bulk counterparts. For example, Vehicle-generated T cells from a DC-depleted culture were less effective than vehicle-generated T cells containing this APC in the culture. This finding indicates that these APCs are, as predicted, important for imparting benefit to transferred T cells. Also, the removal of macrophages from the vehicle culture enhanced this therapy. This finding may indicate that macrophages in the culture, if left unstimulated, actually skew the outgrowth of more suppressed and less functional T cells. Future studies are needed to address how a macrophage population in culture may skew the expanded T cell product.

What remained striking to us was the potent activity of CpG on B cells to generate antitumor CD8^+^ T cells for ACT. However, depleting B cells from the cell culture did not satisfy the question of whether B cells alone were able to generate this potent cell therapy product via CpG. This distinct question is of critical importance, as the translation of this finding would be more feasible if isolated B cells could impart this biology on T cells via CpG without the need for other immune cells. When purified B and CD8^+^ T cells were co-cultured, the addition of CpG improved their activity when infused into mice with large tumors. Of note, B cells slightly enhanced CD8^+^ T cells immunity even when CpG is not present. The finding that B cells were critical for the efficacy of CpG-improved T cell therapy leads to several questions of the mechanism of these cells’ interaction. For example: is the improvement with CpG B cell/T cell contact-dependent, or are the alterations driven by a B cell-secreted factor? Answers to these questions could lead to additional novel strategies to improve cell therapy for cancer.

B cells are emerging as a potentially critical presence in successful immunotherapies. In fact, three exciting studies revealed that the presence of B cells in tertiary lymphoid structures (TLS) in several malignancies served as a powerful prognostic indicator of successful immune checkpoint blockade (ICB) therapies^44–46^. Each of these groups performed in-depth analyses to derive more information about how B cells in the tumor may be contributing to tumor immunity. Several features unique to ICB responders arose: the presence of specific B cells subsets, altered BCR clonality and diversity, and B cell/T cell interactions within the TME. However, whether or how each of these observations drive response to checkpoint blockade and if their presence is predictive of response to ICB in other malignancies remains to be determined. As B cells can play multiple roles in the immune response – antibody production to tumor antigens, presentation of antigens to CD4 and CD8 T cells, and cytokine/chemokine production – it is likely that they contribute to tumor immunity *in vivo* in a number of ways. Indeed, tumor-infiltrating B cells (TIL-B) have been shown to act as APCs *ex vivo* to CD4^+^ TIL from human NSCLC tumors^47^. Thus, exploiting and enhancing this interaction by incorporating TLR agonists, is a logical next step. The findings from these clinical reports alongside our findings prompt another question of whether B cells, *in vivo*, are also critical for the response of adoptive cell therapies for cancer.

Our findings can be directly translated to improve cell therapies. In TIL-based ACT therapies, two approaches could be used: 1) at the onset of culture when many tumor pieces harbor a TIL-B population or 2) during rapid expansion in which T cells are expanded in the presence of PBMC feeder cells to large numbers before returned to the patient. We posit that incorporating CpG into either step of *ex vivo* expansion could lead to improved cellular therapies. Ongoing studies in our lab seek to understand how to best use CpG in clinical trials at various institutions. Collectively, our work demonstrates, for the first time, the importance of B cells in generating potent CD8^+^ T cell products for cancer immunotherapy.

## METHODS

### Mice and Tumor Cell Lines

Pmel-1 TCR transgenic mice were purchased from the Jackson Laboratories and bred in the in-house animal facilities (comparative medicine department) at the Hollings Cancer Center of the Medical University of South Carolina (MUSC). C57BL/6 CD45.1 mice were purchased from the NCI Frederick laboratories for use in *in vivo* tumor experiment studies. Mice used for tumor experiments were between 6-10 weeks old. All animals were housed and underwent experimentation in accordance with MUSC’s Institutional Animal Care and Use Committee (IACUC) and under the supervision and support of the Division of Lab Animal Resources (DLAR). IACUC approval was obtained prior to all animal studies and procedures. We received the B16F10 (H-2^b^) melanoma cell line from Nicholas P. Restifo, which was used in *in vivo* tumor studies.

### T cell Culture

#### Pmel-1 cells

Pmel-1 transgenic T cells were obtained via the culture of whole pmel-1 splenocytes. Pmel-1 splenocytes were seeded at 1e^6^/mL in a 24-well plate and activated with 1μM human gp100 (hgp100) peptide (unless indicated otherwise) in the presence of 100IU/mL IL-2 (NIH repository). At the time of activation (unless otherwise stated) either mouse CpG-ODN 1668 (5’-tccatgacgttcctgatgct-3’) (Class B) was added to the culture at 0.5μg/mL or vehicle control (endotoxin-free water) was added at a matched volume (InvivoGen). On day 2 of culture, half of the culture media was replaced with fresh media with 100IU/mL IL-2 in the new media. From day 3 of culture on cells were split to a concentration of 1e^6^ cells/mL and supplemented with fresh media containing 100IU/mL IL-2. On day 7 of culture, T cell cultures were assayed using flow cytometry and then directly used for *in vivo* tumor experiments where indicated. *CD8^+^ T cell negative isolation:* For experiments in which T cells were purified from the bulk pmel-1 splenocytes before culturing we used the EasySep Mouse CD8^+^ T cell Isolation Kit (Stem Cell, Cat# 19853A) following the manufacturer’s protocol. Prior to cell culture, the T cell isolate was assayed for the purity of the product which was always more than 90% pmel-1 transgenic T cells (identified via expression of the Vβ13 chain of the TCR). *B cell negative isolation:* B cells were negatively isolated using the EasySep Mouse B cell Isolation Kit (Stem Cell, Cat# 19854) following the manufacturer’s instructions. B cell purity was assayed post isolation and prior to cell culture with purified CD8^+^ T cells. In recombination experiments, B cells were cultured with CD8^+^ T cells at a 4:1 B:T cell ratio and activated and expanded in culture as described above. *Immune cell subset depletion:* From the bulk pmel-1 splenocytes we depleted either CD4^+^ T cells, NK cells, dendritic cells, macrophages, or B cells using subset marker-specific PE labeling followed by anti-PE microbead targeting and positive selection via column isolation. The negative fraction was used for culture as it does not contain the positively labeled cell subset. Briefly, pmel-1 splenocytes are labeled with PE-conjugated antibodies: either αCD4-PE, αNK1.1-PE, αCD11c-PE, αF4/80-PE, or αCD19-PE (antibodies listed in **Supplemental Materials Table 1**) to mark CD4^+^ T cells, NK cells, dendritic cells, macrophages, or B cells respectively. Cell mixtures were then further labeled with Anti-PE MicroBeads from MACs Miltenyi Biotec (130-048-801) and then underwent magnetic separation with the MACs LS Columns (130-042-401) according to the manufacturer’s instructions. Both positive and negative fractions were collected into 15mL tubes, and assayed via flow cytometry for the presence of the PE-conjugated cell specific marker as well as another secondary marker, when applicable, to identify the presence of each subset in each fraction. Percent depletions of each subset were as follows: 97.7, 92.5, 99.6, 99.5, and 69.1 of F4/80+ macrophages, CD11c^+^ DCs, CD19^+^ B cells, CD4^+^ T cells, and NK1.1^+^ NK cells. The negative fraction was cultured as described above, using hgp100 peptide activation, 100IU/mL IL-2, vehicle or CpG treatment on day 0. Bulk pmel-1 splenocytes were cultured in the same manner as controls.

#### Human OCSCC TIL culture (oral cavity)

Our TIL culture method was adapted from the method by Dudley et al^48^. Briefly, tumor specimen was collected fresh from a patient with oral cavity squamous cell carcinoma (OCSCC) and cut into 1-3mm^2^ fragments before seeding in complete media (RPMI) supplemented with 6000U/mL recombinant IL-2 (NCI repository). At the time of seeding 3 tumor fragments were additionally treated with either vehicle control (endotoxin-free water) or CpG ODN 2006 (InvivoGen) at 0.5ug/mL. TIL was left untouched for 5 days in culture to allow cell egress from the tumor piece. After 5 days 1mL/well (24-well plate) was removed without disturbing the cells settled on the bottom of the well and replaced with fresh media with 6000U/mL IL-2. Cells are monitored and media is replaced every 3-5 days. Once cells become confluent in the well, the tumor piece is removed and cells are re-plated and maintained at 1-1.5× 10^6^ cells/mL in a 24 well plate.

### Adoptive Cell Therapy

B16F10 cells from culture were washed twice, resuspended in sterile PBS, and injected subcutaneously to the abdomen at 5e^5^ cells/mouse (in 200uL). Tumors were established for 5 days *in vivo* prior to ACT and mice were irradiated with 5Gy total body irradiation (TBI) the day before ACT. Prior to treatment, mice were randomized according to tumor size to distribute tumor size evenly among groups. If no tumor was detectable four days post tumor cell injection, mice were not included in the study. 4-5e^6^ pmel-1 cells expanded one week *in vitro* were resuspended in sterile PBS and transferred via tail vein injection. Starting from day 0 of ACT, tumors were measured 2x per week with handheld calipers by a lab member blinded to the treatment group until tumors reached protocol endpoint (400mm^2^). Mice euthanized prior to endpoint for tissue collection and biodistribution analysis were identified and allocated prior to commencing treatment.

### Mouse Blood and Tissue Collection

#### Peripheral blood

Mouse peripheral blood was collected at the indicated time points post-ACT from the mandibular vein into 1.5mL Eppendorf tubes containing 0.125 M EDTA. All blood was pelleted, and red blood cells were lysed (Biolegend). Immune cells were then assayed via flow cytometry. *Tumor collection:* Tumors were excised and placed in culture media on ice until tissue dissociation. Tumors were dissociated into a single cell suspension via mashing and washing over a 70uM screen and collected into 50mL tubes. Cell suspensions were further filtered through a 35uM filter prior to flow cytometry analysis to prevent clumping.

### Proteomics

#### Sample preparation

Cells were lysed in 9M urea, 50 mM Tris pH 8, with 100 units/mL Pierce Universal Nuclease (ThermoScientific) added. The concentration of protein was measured using the BCA assay (ThermoScientific). Fifty micrograms (μg) of protein from each sample were brought to the same final volume using the lysis buffer and reduced in 1 mM 1,4-Dithiothreitol (DTT), and alkylated in 5.5 mM iodoacetamide (both from Thermo Scientific). The urea concentration was then reduced to 1.6M using 50 mM ammonium bicarbonate. The samples were digested with Lys-C (Waco) at a 1:50 protease: protein ratio for 3 hrs. at room temperature followed by trypsin (Sigma) at 37°C for 18 hours at a 1:50 protease: protein ratio. The resulting peptides were desalted using C18 Stage Tips (ThermoScientific) using a standard protocol. The elution from the stage tip for each sample was dried in a Speed Vac and stored at −80°C.

#### Liquid Chromatography and Mass Spectrometry Data Acquisition Parameters

Dried peptides were dissolved in 15 μL of 2% acetonitrile (ACN)/ 0.2% formic acid (FA) and 5 μL of this injected. Peptides were separated and analyzed on an EASY nLC 1200 System (ThermoScientific) in-line with the Orbitrap Fusion Lumos Tribrid mass spectrometer (ThermoScientific) with instrument control software v.4.2.28.14. Peptides were pressure loaded at 1,180 bar, and separated on a C18 reversed phase column (Acclaim PepMap RSLC, 75 μm x 50 cm (C18, 2 μm, 100 Å)) (ThermoFisher) using a gradient of 2% to 35% B in 180 min (Solvent A: 0.1% FA, 2% ACN; Solvent B: 80% ACN/ 0.1% FA) at a flow rate of 300 nL/min. The column was thermostated at 45 °C. Mass spectra were acquired in data-dependent mode with a high resolution (60,000) FTMS survey scan, mass range of m/z 375-1575, followed by tandem mass spectra (MS/MS) of the most intense precursors with a cycle time of 3 s. The automatic gain control target value was 4.0e5 for the survey scan. Fragmentation was performed with a precursor isolation window of 1.6 m/z, a maximum injection time of 50 ms, and HCD collision energy of 35%; the fragments were detected in the Orbitrap at a 15,000 resolution. Monoisotopic-precursor selection was set to “peptide”. Apex detection was not enabled. Precursors were dynamically excluded from resequencing for 15 sec with a mass tolerance of 10 ppm. Advanced peak determination was not enabled. Precursor ions with charge states that were undetermined, 1, or > 7 were excluded from fragmentation.

#### Mass Spectrometry Data Processing

Protein identification and quantification were extracted from raw LC-MS/MS data using the MaxQuant platform v.1.6.3.3 with the Andromeda database searching algorithm and label-free quantification (LFQ) algorithm^49–51^. Data were searched against a mice Uniprot reference database UP000000589 with 54,425 proteins (April, 2019) and a database of common contaminants. The false discovery rate (FDR), determined using a reversed database strategy, was set at 1% at the protein and peptide level. Fully tryptic peptides with a minimum of 7 residues were required including cleavage between lysine and proline. Two missed cleavages were permitted. The LFQ feature was on with “Match between runs” enabled for those features that had spectra in at least one of the runs. The “stabilize large ratios” feature was enabled, and “fast LFQ” was disabled. The first search was performed with a 20ppm mass tolerance; after recalibration, a 4.5 ppm tolerance was used for the main search. A minimum ratio count of 2 was required for LFQ protein quantification with at least one unique peptide. Parameters included static modification of cysteine with carbamidomethyl and variable N-terminal acetylation and oxidation of methionine. The protein groups text file from the MaxQuant search results was processed in Perseus v. 1.6.8.0^52^. Identified proteins were filtered to remove proteins only identified by a modified peptide, matches to the reversed database, and potential contaminants. The normalized LFQ intensities for each biological replicate were log2 transformed. Quantitative measurements were required in at least two out of three biological replicates in at least one of the treatment groups. Gene ontology and Reactome pathway annotations were added from the murine Uniprot reference proteome. The mass spectrometry proteomics data have been deposited to the ProteomeXchange Consortium via the PRIDE^53^ partner repository with the dataset identifier PXD022909. PCA analysis was performed in Perseus. A volcano plot was generated in Perseus using a signifcance threshold of a two sided t test adjusted pvalue (<0.05 FDR) and a S0 parameter of 0.1. Protein set enrichment analysis was performed using the ToppFun function of the ToppGene Suite (https://toppgene.cchmc.org/enrichment.jsp), where the proteins increased significantly (two sided t test adjusted pvalue (<0.05 FDR) and a S0 parameter of 0.1) were assed.

### Flow Cytometry

Flow cytometry was performed on either a BD FACSVerse or BD LSRFortessa X-20 instrument and subsequently analyzed using FlowJo software (BD). For extracellular staining, cells were suspended in FACS Buffer (PBS + 2% FBS) and incubated with antibodies against cell surface markers (indicated in supplemental Table 1) and Zombie Aqua viability dye for 20 minutes at room temperature or 30 min at 4° C. Cells were then washed with FACS Buffer prior to running on either flow cytometer.

### ImmGen Database Query

Murine Tlr9 transcript expression was assessed in multiple immune cell types using the online database of the Immunological Genome Project (https://www.immgen.org/). We queried the ImmGen ULI RNASeq database under the Gene Skyline databrowser.

### Statistics

Graphs and statistical analyses were created and performed in GraphPad Prism 7 or Perseus for proteomics analyses. Survival was compared using a Log-rank (Mantel-Cox) test. To compare two treatment groups, we used either a Two-sample T-test or a Mann-Whitney U test as indicated in figure legends. P values less than 0.05 were considered significant. In average tumor curves each point signifies the average tumor area at that time and the error bars indicate standard error of the mean. In dot-plots of replicate analyses mean and standard deviation are indicated. To identify significantly regulated proteins a Student t-Test was performed with a permutation-based FDR cutoff of 0.05 and S0 = 0.1^54^.

### Study Approval

All animal procedures were approved by the Institutional Animal Care & Use Committee of the Medical University of South Carolina, protocol number 0488.

**Table 1.**

| **Target**<br>**(anti-mouse)** | **Clone** | **Fluorophore** | **Manufacturer** | **Catalogue**<br>**number** |
| --- | --- | --- | --- | --- |
| CD8 | 53-6.7 | PerCpCy5.5 | Biolegend | 100734 |
|  | 53-6.7 | PE | BD<br>Biosciences | 553033 |
| CD4 | GK1.5 | APC Cy7 | BD<br>Biosciences | 552051 |
| Vβ13 | MR12-3 | APC | BD<br>Biosciences | 561542 |
| B220 | RA3-62B | APC | Invitrogen | 17-0452-82 |
| MHCII | M5/114.15.2 | PE | Biolegend | 107607 |
| CD11c | N418 | BV41 | Biolegend | 117343 |
|  | HL3 | PE | BD<br>Biosciences | 557401 |
| F4/80 | BM8 | PeCy7 | Biolegend | 123113 |
|  | T45-2342 | PE | BD<br>Biosciences | 565410 |
| NK1.1 | PK136 | FITC | BD<br>Biosciences | 553164 |
|  | PK136 | PE | BD<br>Biosciences | 557391 |
| CD19 | 1D3 | PE | BD<br>Biosciences | 553786 |
| IL-2Ralpha | 7D4 | FITC | BD<br>Biosciences | 553072 |
|  | PC61 | PE | BD<br>Biosciences | 553866 |
| ICOS | C398.4A | BV421 | Biolegend | 313524 |
| CD39 | Duha59 | Pe-Cy7 | Biolegend | 143806 |
| CD44 | IM7 | PerCp-Cy5.5 | Biolegend | 103032 |
| TLR9 | M9.D6 | FITC | Invitrogen | 11-9093-80 |
| CD3 | 17A2 | eflour450 | eBioscience | 48-0032-82 |
| CD64 | X54-517.1 | Pe-Cy7 | Biolegend | 139313 |
| **Target**<br>**(anti-human)** | **Clone** | **Fluorophore** | **Manufacturer** | **Catalogue**<br>**number** |
| CD8 | RPA-T8 | PerCp-Cy5.5 | Biolegend | 301032 |
| CD4 | RPA-T4 | PE | eBioscience | 12-0049-42 |
| ICOS | 4B4-1 | V450 | Biolegend | 309819 |
| IL-2Ralpha | BC96 | APC-Cy7 | Biolegend | 302614 |
| CD39 | A1 | PE-Cy7 | Biolegend | 328212 |

## AUTHOR CONTRIBUTIONS

**A.S.S.:** Conceptualization, data curation, formal analysis, investigation, methodology, visualization, writing - original draft, writing – review and editing. **H.M.K.:** Data curation, investigation, validation, writing – review and editing. **M.M.W.:** Conceptualization, data curation, investigation, methodology, supervision, writing – review and editing. **C.J.D.:** Data curation, investigation, writing – review and editing. **G. O. R. R.:** Data curation, investigation, writing – review and editing. **A.M.R.R.:** Data curation, investigation, writing – review and editing. **D.M.N.:** Resources, supervision, writing – review and editing. **J.E.T.:** Investigation, methodology, resources, supervision, writing – review and editing. **E.B.:** Investigation, methodology, supervision, writing – review and editing. **M.P.R.:** Investigation, supervision, writing – review and editing. **B.L.:** Investigation, methodology, supervision, writing – review and editing. **C.M.P.:** Conceptualization, data curation, formal analysis, funding acquisition, investigation, project administration, resources, supervision, visualization, writing - original draft, writing – review and editing.

## ACKNOWLEDGEMENTS

This work was supported by NCI F31 CA232646-01A1 and Hollings Cancer Center Graduate Fellowship (A.S.S.), NCI F30 CA243307, NIH T32 GM008716 and DE017551 (H.M.K.), NIH R50 CA233186 (M.M.W.), NIH T32 GM008716, 2019 MUSC CGS Provost Scholarship (G.O.R.R.), NIH T32 AI132164-01 (C.J.D.), NIDCR K08 DE26542 and SCTR UL1TR001450 (D.M.N.), ACS RSG-17-047-01-MPC, NIH NCI R01CA194090 and NIH NIAID R21AI142387 (E.B.), NCI R01 CA222817 (M.P.R.) NIH NCI (R01: CA193939) and NIH NIAID (U01: AI125859) (B.L.), Hollings Cancer Center Proteomics Pilot Award (A.S.S. and C.M.P.), NCI R01 CA175061, R01 CA208514 (C.M.P.), and S10 OD010731 (Orbitrap Fusion Lumos MS, L.E.B), GM103542 (Proteomics Core, L.E.B.) from the National Institutes of Health. We acknowledge technical assistance from the Mass Spectrometry Facility, a University Research Resource, specifically Susana Comte-Walters, Jennifer R. Bethard, Baylye Burnette, and Lauren Ball. We thank Kent Armeson for his guidance on statistical analyses. We thank our collaborators for their valuable feedback: Arman Aksoy, Pinar Aksoy, Elinor Gottschalk, Dimitrios Arhontoulis, Stephen Iwanowycz, Katie Hurst, Brandon Ware, Gregory Lesinski, Ragini Kudchadkar, and David Lawson.

## Notes

### Competing Interest Statement

The authors have declared no competing interest.

