## Supplemental Figures for "B cells imprint adoptively transferred CD8^+^ T cells with enhanced tumor immunity"

**Supplemental Material**

**Supplemental Figure 1. Mouse T cells express nominal TLR9.** A) Murine *TLR9* transcript expression by RNAseq was queried using an online database of the Immunological Genome Project (immgen.org), ImmGen ULI RNASeq, under the Gene Skyline databrowser. Various immune and non-immune cell types are shown with their expression of Tlr9 where expression value ranges are categorized as 'trace' (blue), 'very low' (green), 'low' (yellow), and 'medium' (orange). B) Intracellular expression of TLR9 at baseline in various pmel-1 immune cell types by intracellular flow cytometry (representative histograms on left and biological replicates on right).

**Supplemental Figure 2. CpG does not confer purified T cells with an altered phenotype or enhanced anti-tumor ability** A) Cell counts over time during culture of purified T cells activated and expanded with Vehicle or CpG. B) Expression of surface markers on day 7 of cell culture of purified $CD8^+$ pmel-1 treated on day 0 with vehicle or CpG. C) Tumor area over time of mice treated with NT, vehicle treated purified pmel-1, or CpG treated purified pmel-1. D) Percentage of donor cells (Vb13+CD8+) in the blood of mice from D on D4, 12 and 25 post ACT. Statistics: B) Unpaired T-test, D) Mann-Whitney test. ns, not significant, *p< 0.05.

**Supplemental Figure 3. The signature CpG phenotype and enhanced anti-tumor efficacy do not rely on $CD4^+$ T cells or NK cells.** A) Representative column depletion efficacy on day 0, prior to plating. B) Representative histograms showing expression of a given marker on D7 of cell culture. C) Tumor area over time of mice treated with NT, or Veh or CpG-generated T cell products from CD4 or NK cell-depleted cultures. D) Survival of mice in C. E) Percentage of donor cells in the blood of mice treated in C over time post-ACT. Statistics: D) Log-rank test. E) Mann-Whitney U at each time point. ns, not significant, *p< 0.05, **p< 0.01, ***p< 0.001, ****p< 0.0001.

**Supplemental Figure 4. Co-cultured B and T cells induce the signature phenotype with CpG.** A) Expression of phenotypic markers on day 7 of cell culture from Bulk or purified B + T cell cultures treated with CpG or Vehicle control on Day 0 (3 biological replicates). Statistics Unpaired T-tests *$p < 0.05$, **$p < 0.01$, ***$p < 0.001$, ****$p < 0.0001$.

### Supplemental Figure 1. Mouse T cells express nominal TLR9

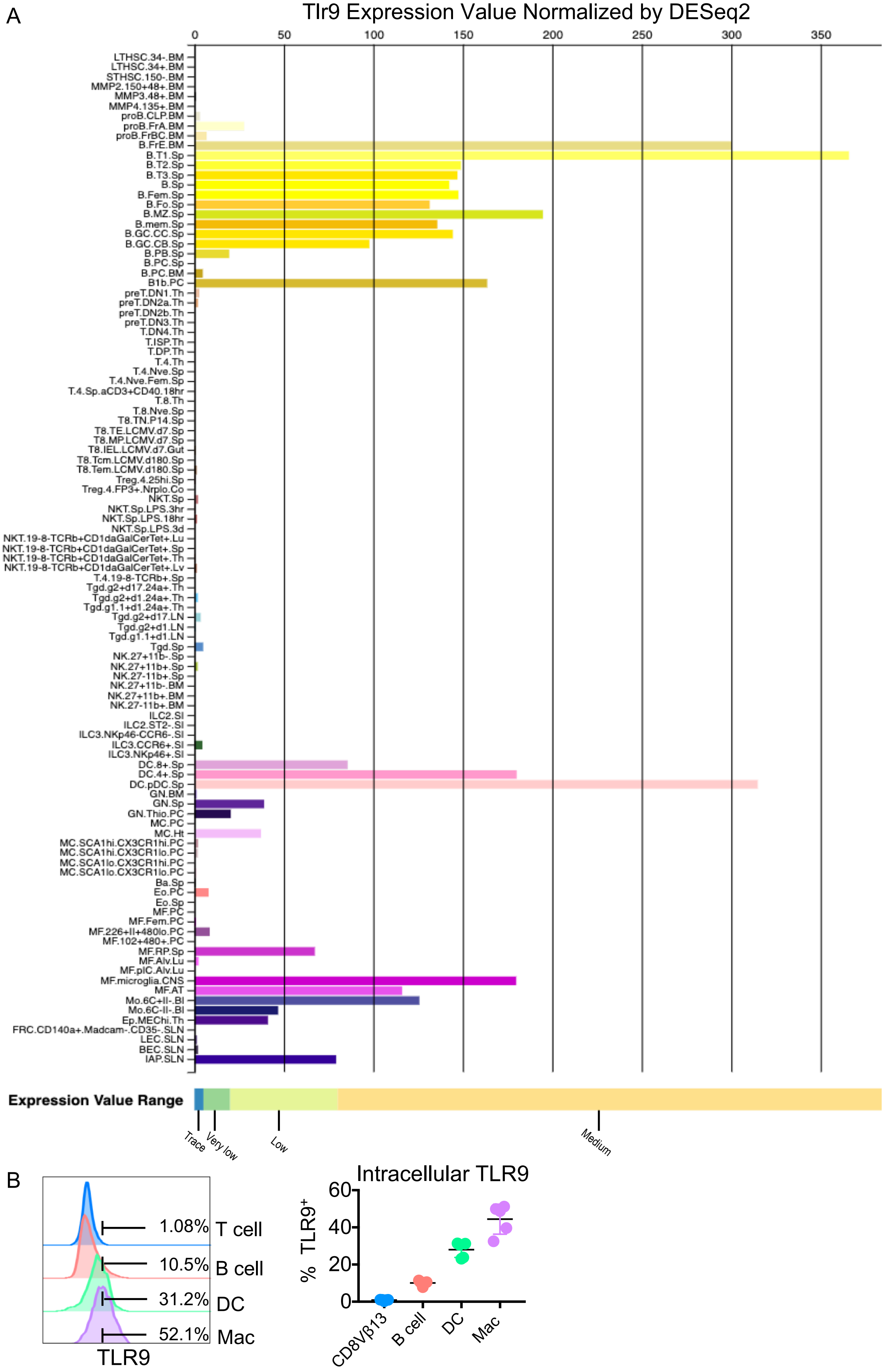

**Supplemental Figure 2. CpG does not confer purified T cells with an altered phenotype or enhanced anti-tumor ability**

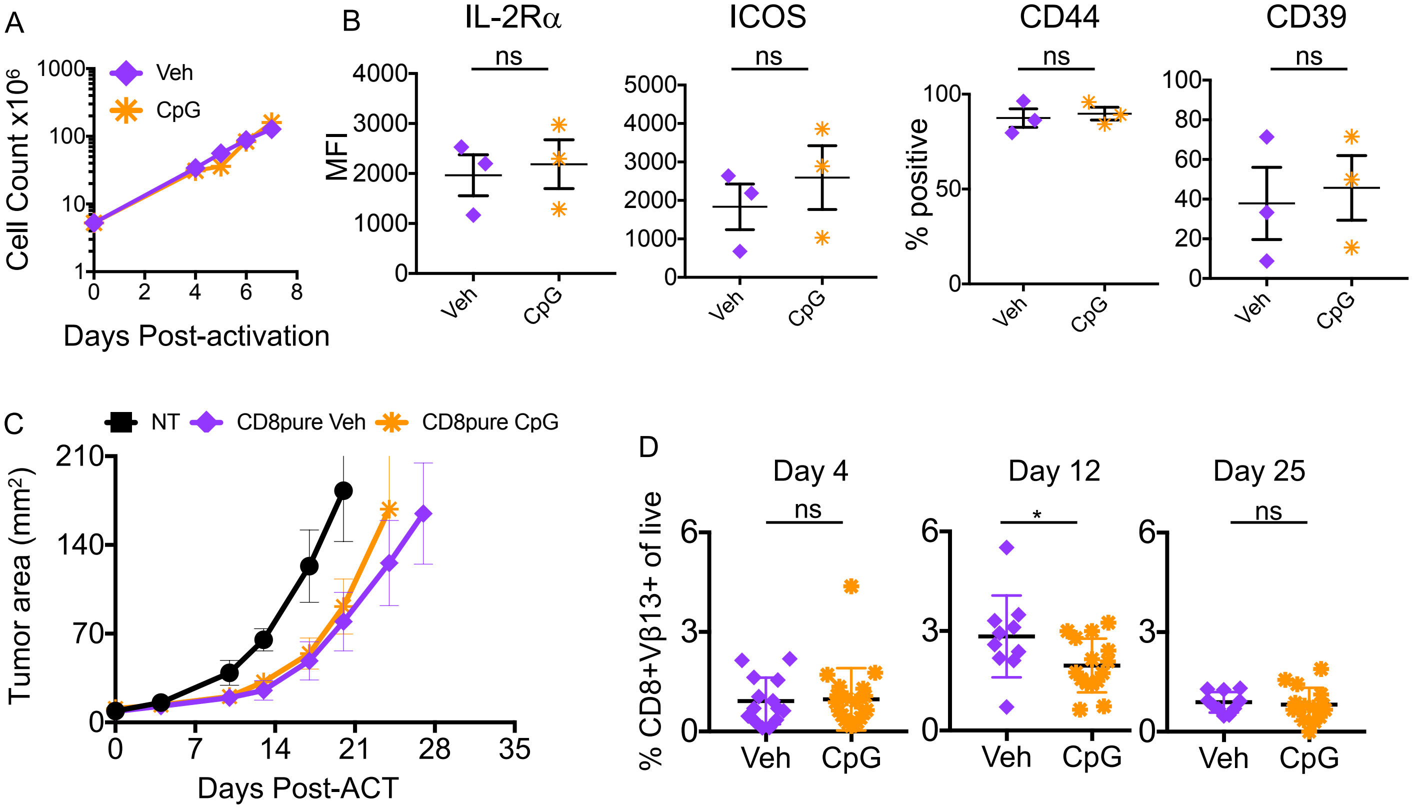

**Supplemental Figure 3. The signature CpG phenotype and enhanced anti-tumor efficacy do not rely on CD4⁺ T cells or NK cells.**

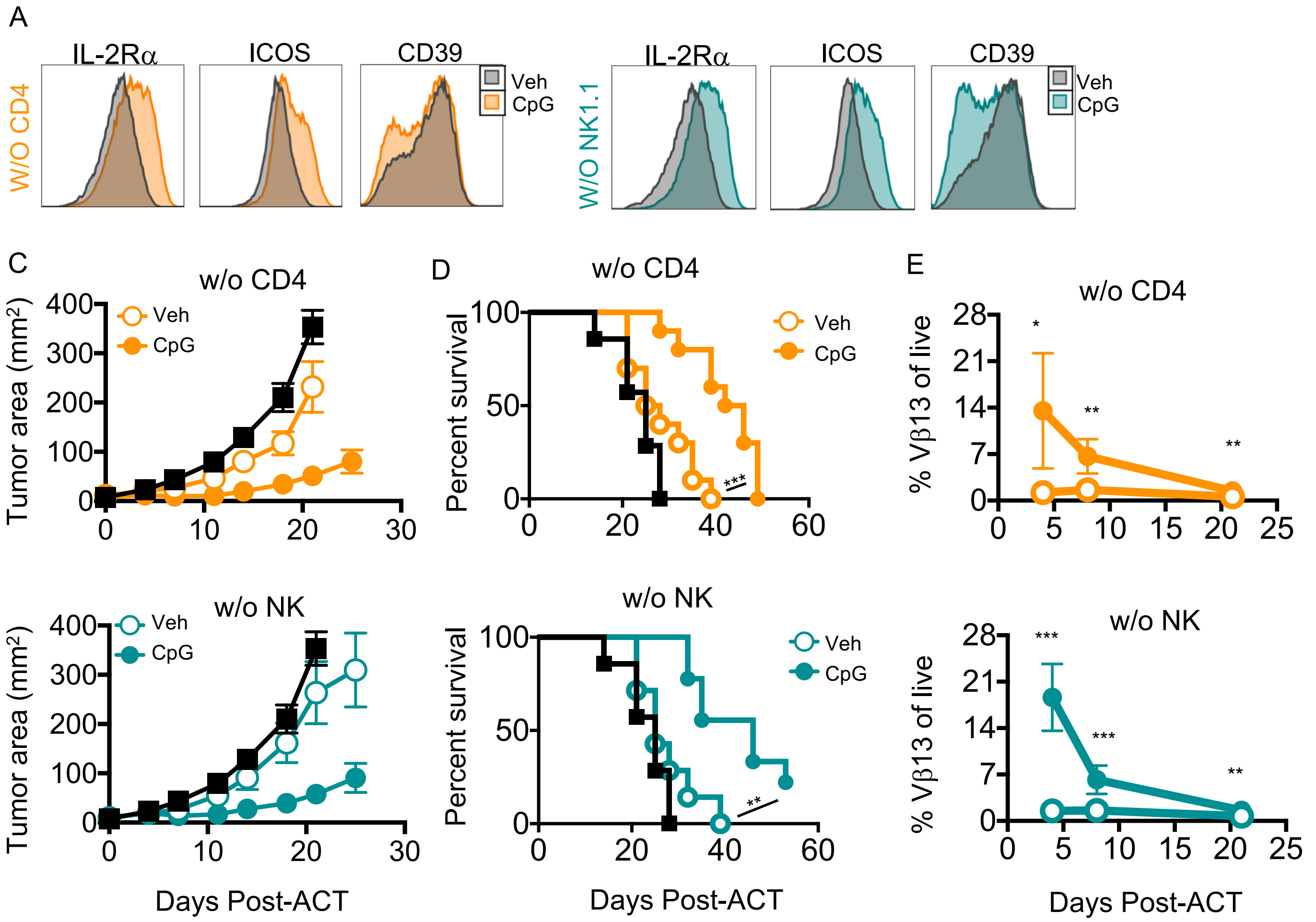

**Supplemental Figure 4. Co-cultured B and T cells induce the signature phenotype with CpG**

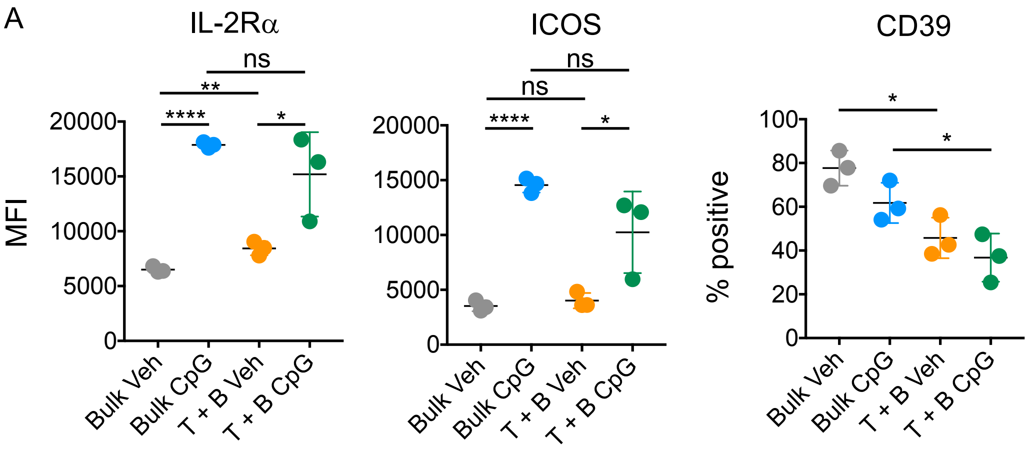
